# *In silico* comparison of novel proteins from Plasmodium Falciparum as candidate drug targets against malaria

**DOI:** 10.1101/2024.12.13.627476

**Authors:** Angela Merici Giannetta Surya, Felicia Austin, Arli Aditya Parikesit

**Affiliations:** Bioinformatics Department, School of Life Science, Indonesia International Institute for Life Sciences, Jakarta 13210, Indonesia

**Keywords:** *Plasmodium Falciparum*, anti-malarial drug target, novel protein drug target, proteome mining, protein-protein interactions

## Abstract

Malaria is a parasitic disease with a persistent prevalence in multiple parts of the world. *Plasmodium falciparum*, the deadliest human-infecting strain, has been a rising concern as the multi-faceted life-cycle poses a challenge in developing anti-malarial treatments while the effectiveness of present drugs begins to decline. As such, various research has been done to identify new drug targets leveraging modern computational biology techniques, such as proteomics. The objective of this study is to evaluate potential drug targets by comparing novel proteins in *P. falciparum* as reported in recent literature (2018-2022) based on their structural analysis of their 3D models along with the druggability properties and protein-protein interactions. A list of 22 proteins were acquired from 48 literature, in which only 7 can be considered as a favorable candidate protein drug target due to their non-human homologous nature, ERRAT score, residues percentage in favourable spot in Ramachandran plot, and high druggability score. Following an examination of protein-protein interactions, it is found that among the evaluated proteins only PF3D7_1438600 exhibits a sufficiently robust network, highlighting it to be the most promising candidate for drug targeting. This study demonstrates the valuable role of proteomics in uncovering potential drug targets and highlights its significance in driving emerging research efforts.

## Introduction

The vector-borne parasitic infection malaria is a severe worldwide health issue that accounts for widespread mortality across the globe, especially low- and middle-income countries (LMIC). Reported by the Word Health Organization (2022), 247 million malaria cases were reported to be found in 84 malaria endemic countries—mostly sub-Saharan Africa countries—and an estimated 619,000 deaths were caused by malaria in 2021 alone [30]. The infection is caused by the parasite *Plasmodium* sp. through the bite of a female Anopheles mosquito, infecting the host’s red blood cells through the liver and causing life-threatening cell and tissue damage [10]. There are currently six known human-infecting *Plasmodium* species: *Plasmodium falciparum, Plasmodium vivax, Plasmodium malariae, Plasmodium ovale curtisi, P. ovale wallikeri* and *Plasmodium knowlesi*, with *P. falciparum* known to be the most dangerous and fatal [18].

Despite the multiple global efforts and malaria management programs, the eradication of the disease is far from over. *P. falciparum* is found to have a multifaceted life-cycle with diverse variations of stage-specific antigens at each phase, presenting a formidable challenge in developing effective strategies that can induce a generalized intervention targeting these antigens [3]. Another arising challenge was the introduction of mutated drug-resistant *P. falciparum*. It has been documented that many approved and widely used anti-malarial drugs are losing their efficacy as *P. falciparum* gains resistance overtime, notably chloroquine—the most common anti-malarial drug, is reported to be no longer effective in almost the majority of the world population [27]. This resistance was reported as long as decades ago, and has started to emerge since the 1950s, so the polymorphisms *P. falciparum* has undergone presents a significant challenge towards the malaria elimination program.

The use of proteomics approach towards the discovery of drug targets has been deemed as important [22]. Proteomics have been utilized to study the protein functions and its changes throughout a disease in order to determine which proteins are specifically changed because of the disease and what it does. Various studies have explored the possibilities of different proteins from the *P. falciparum* as novel drug targets for malaria [2, 25]. Even so, those studies have come up with different results with different methodologies which meant that the effectiveness of those novel drug targets have not been compared as of yet. This study aims to obtain the best potential anti-malarial drug targets from a list of recently discovered novel *P. falciparum* proteins.

## Materials and Methods

The pipeline for this research was adapted from a malaria novel drug target study by Ali et al. (2021) [2]. Adjustment was done as the source to gather the sample for this research differs from the referenced study.

**Figure 1.**
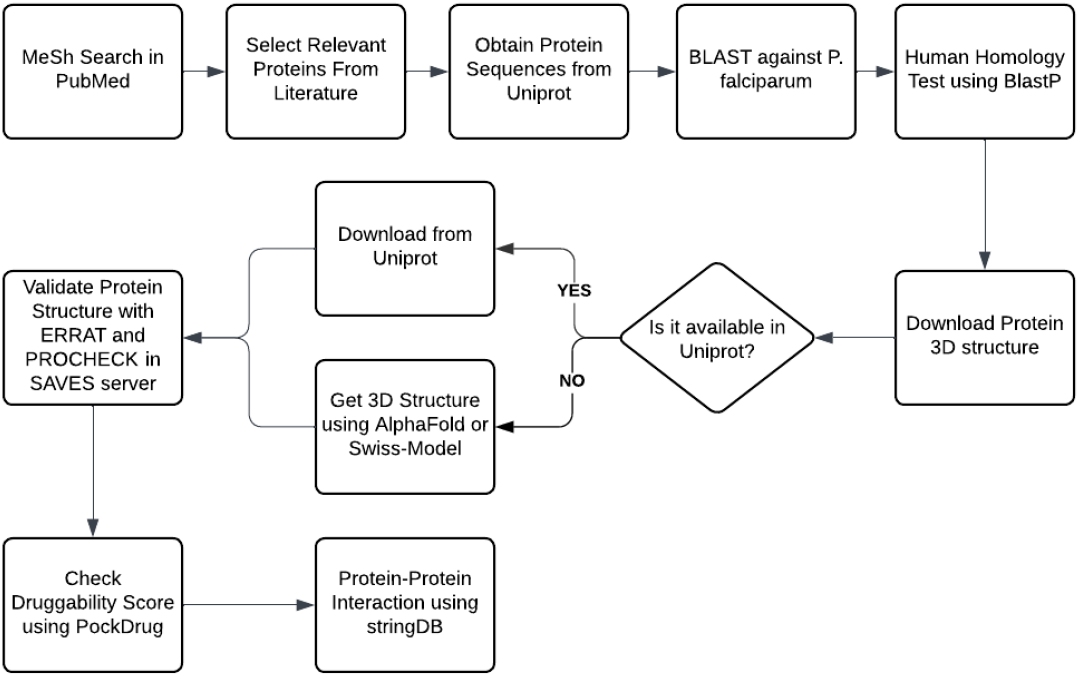
Workflow for determining the best candidate drug target

### Protein gathering

The protein list was gathered using NCBI PubMed from other literature published since 2018 which contains the keyword plasmodium falciparum, anti-malarial, protozoan proteins, and novel drug targets. In total, there were 48 research articles which were then filtered with criteria of i) research discovered new protein target, ii) research listed the protein researched, and iii) research that has not been confirmed in vitro/vivo. The filter resulted in 4 papers as sources and a total of 22 proteins as novel drug targets. The sequence of the proteins obtained was then inputted through BLAST in order to confirm the existence of the protein in *P. falciparum* [7]. Afterwards, the sequence was run through BlastP to examine its homology against human proteomes. The protein must be non-homolog to the human host proteome as to avoid the drug to target the host instead. By the end of this stage, only 11 proteins were confirmed as a *P. falciparum* and a non-human homolog protein.

### Druggability score

The 3D conformer of each protein was gathered using the Uniprot database in the form of a PDB file [29]. If not available, the sequence was then run through AlphaFold in order to predict its 3D structure or through Swiss-Model [6, 16]. To confirm the quality of the 3D model, the result was then filtered with the parameter of ERRAT quality factor ≥ 90 and residues in favored region of Ramachandran plot ≥ 80% to be considered as a good model using the SAVES server [9, 19]. There were only 7 proteins left that passed the good quality model. In order to determine if the protein were able to be classified as a drug target, druggability score was then obtained using the PDB file through the PockDrug web server [15]. The protein pocket would be considered as druggable if the druggability score is > 0.5. Each protein network was then obtained using the STRING database with the parameter of *Plasmodium falciparum* as organisms, full STRING network as network type, high confidence (0.700) as required score, and no more than 10 interactors as size cutoff [26].

## Results and Discussion

Vital cellular functions necessary for pathogens survival are regulated by protein-coding genes [24]. 30% of the annotated 5,389 proteins from the whole proteome of *P. Falciparum* is still denoted as hypothetical proteins [25]. Previous studies have focused on identifying potential targets for new anti-malarial drugs by assigning functions and determining the structures of uncharacterized proteins through protein annotation. Over the course of the recent five years, we identified a number of 22 newly-annotated proteins that were investigated as novel anti-malarial drug targets, as listed in Table 1 [23, 2, 25, 31]. Further analysis of all 22 proteins are found to be specific to *P. falciparum*. The protein sequences obtained were compared to the human proteome from the NCBI RefSeq database using BLASTp. Proteins that showed query coverage and percent identity of 35% or higher were excluded. As a result, a total of 11 non-homologous proteins were identified (Table 2). The structure of these proteins were acquired from the Uniprot database, and only one protein (PF3D7_0724700) structure was found unavailable, and therefore predicted using the Swiss-model [6]. The predicted structure was acquired from an uncharacterized protein (PGSY75_0724700) belonging to *Plasmodium gaboni* and BLAST results against the Uniprot database showed 85.5% similarity in the protein sequence. Validation of the protein structures are done in SAVES v6.0 ERRAT and PROCHECK features (Table 2). Proteins with overall quality factor > 90 from ERRAT and > 80% protein residues found in the favoured regions in the Ramachandran plot are validated as a high quality model [21]. Finally, a selection of seven proteins was identified as a priority for further analysis of their druggability. These proteins are hypothesized to possess potential drug binding sites and can play essential roles as central hub proteins within the biological network of P. falciparum.

**Table 1.**
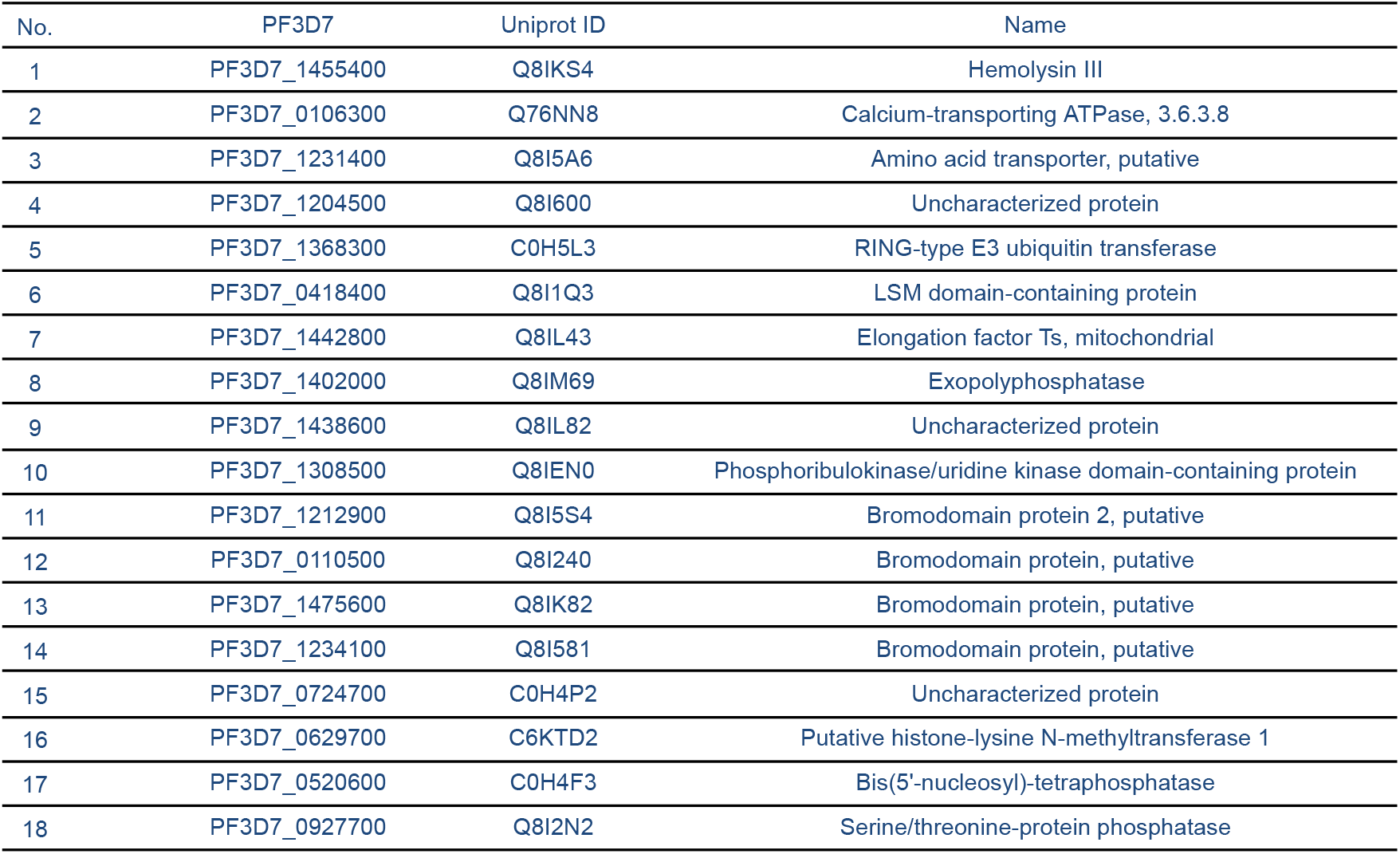
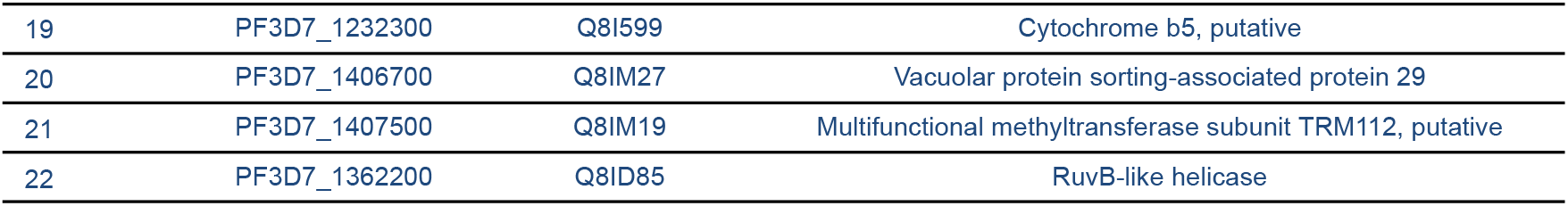
The list of proteins gathered from various literature.

**Table 2.** The list of proteins that are confirmed to be *P. falciparum* and non-human homolog.

| No. | PF3D7 | Name | ERRAT | Ramachandran (%) |
| --- | --- | --- | --- | --- |
| 1 | PF3D7_1455400* | Hemolysin III | 99.2701 | 94.3 |
| 2 | PF3D7_1231400 | Amino acid transporter, putative | 88.4763 | 66 |
| 3 | PF3D7_1204500* | Uncharacterized protein | 97.6 | 88.7 |
| 4 | PF3D7_1368300* | RING-type E3 ubiquitin transferase | 98.0769 | 88.9 |
| 5 | PF3D7_0418400* | LSM domain-containing protein | 100 | 91 |
| 6 | PF3D7_1442800 | Elongation factor Ts, mitochondrial | 82.7381 | 81.1 |
| 7 | PF3D7_1402000* | Exopolyphosphatase | 96.2422 | 91.6 |
| 8 | PF3D7_1438600* | Uncharacterized protein | 99.2565 | 90.6 |
| 9 | PF3D7_1308500* | Phosphoribulokinase/uridine kinase domain-containing protein | 94.4785 | 88.2 |
| 10 | PF3D7_0110500 | Bromodomain protein, putative | 89.4191 | 94.7 |
| 11 | PF3D7_0724700 | Uncharacterized protein | 85.1455 | 75.3 |

\*Signifies the proteins that pass the ERRAT and Ramachandran score

### Druggability Analysis

The term “druggable” target refers to a protein with the ability to interact with a drug compound and the interaction initiates a specific biological response characterized to the disease progression [28]. When interacting with the lead compounds, it is expected that the desired therapeutic effect of the drug compound is elicited, such as function alteration of signaling molecules, or inhibition of an enzyme activity, that might have contributed to the development of the disease. Druggability analysis can be done in silico [11]. Identifying and focusing on druggable targets increases the likelihood of successfully developing drugs that can interact with these targets and modulate disease-related processes.

Assessing the druggability of a target can involve examining the presence of protein “pockets,” which are cavity-like regions within a protein that can accommodate small molecules. These pockets provide potential binding sites where small molecules can interact with the protein. One way to evaluate druggability is by determining if these pockets contain suitable surface binding sites that possess pharmacophoric features [28]. To identify the binding sites and determine its druggability, we use PockDrug, which uses combinations of geometry, hydrophobicity, and aromaticity to evaluate pocket properties impacting druggability [8].

The other variables listed in table 3 were used in calculation of the druggability score in PockDrug. Vol. Hull counted the volume of the convex hull which means the area of the pocket, while Hydroph. Kyte calculated the hydrophobicity based on the properties of the residues. Polar Res. and Aromatic Res calculate the frequency of polar residues and aromatic residues respectively. Otyr atom calculates the presence of a hydroxyl group in the tyrosine side chain which has a big role in binding drug molecules, while Nb. Res. states for the number of pocket residues.

**Table 3.**
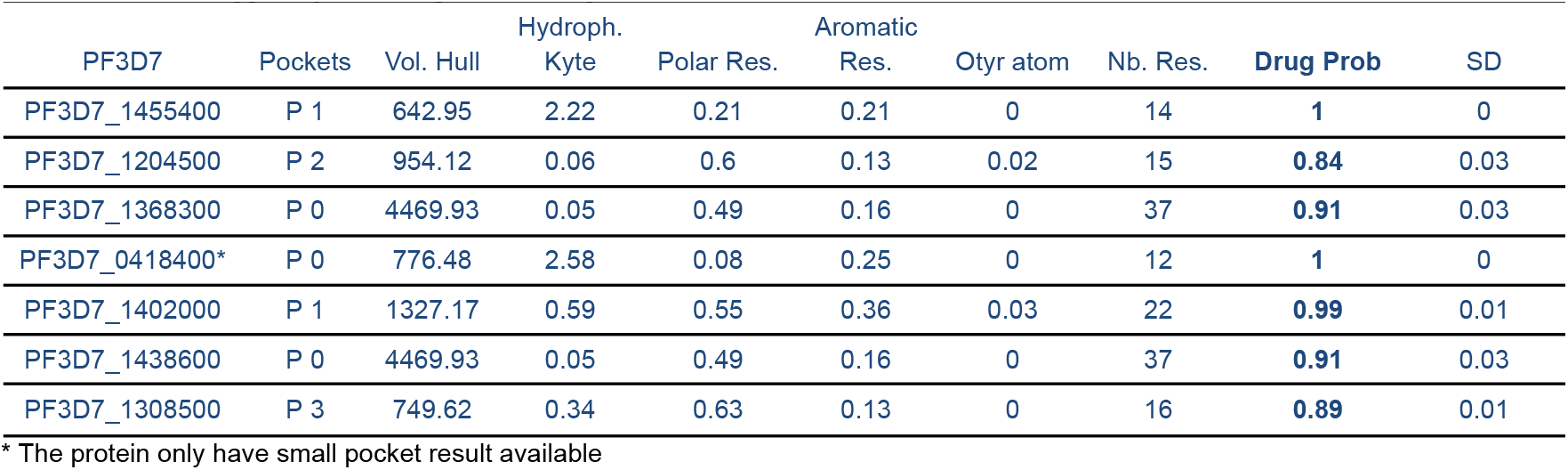
Protein druggability according to PockDrug.

All of the proteins could be considered as a druggable protein as their druggability scores are all >0.5, with the range of (0.84-1). The proteins with the best drug probability score are tied between PF3D7_1455400 and PF3D7_0418400 with both having a score of 1. However it is to be noted that albeit the score of 1, PF3D7_0418400 only has a small pocket which was considered by the pocket having less than 14 residues. A smaller pocket has its own limitations which include a more limited amount of ligand that can bind to the pocket [15]. Thus, it means that the protein PF3D7_0418400 while having a good probability as a potential target protein for malaria, the protein PF3D7_1455400 is still a better candidate due to having a wider range of ligands (potential drugs) to be binded with.

### Protein-protein Interactions (PPIs)

From the list of proteins before, there were only two proteins which have a protein-protein interaction (PPI) network which is the PF3D7_1438600 and PF3D7_1455400. Between the two proteins, PF3D7_1455400 results are unfavorable as it only has 2 average node degrees when compared to PF3D7_1438600 that have the average node degree of 4.7. The score for the ten interactors with PF3D7_1438600 ranges from 0.725 to 0.999, therefore the confidence score of the interaction is quite high. From the PF3D7_1438600 PPI, the protein PF10_0114 plays an important role in *P. falciparum* as it functions in nucleotide excision repair and modulation of proteasomal degradation [17]. By targeting the PF3D7_1438600 protein, there might be a chance of disrupting the DNA repair process of the *P. falciparum* thus potentially killing the protozoa eventually. Another protein called MAL8P1.51 plays a role in translocating protein in the endoplasmic reticulum (ER) [26]. The protein PFD0725c is an ATPase protein which plays a role in delivering tail-anchored (TA) proteins to the ER at the post-translational stage. Research on PfSec22 in *P. falciparum* revealed that this protein displays unique trafficking patterns within the parasite [4]. PfSec22 is found to be orthologous to Sec22, found in the ER-Golgi intermediate compartments (ERGIC), where it facilitates efficient membrane fusion and helps assemble larger protein complexes through its ability to form homodimers [1]. This presents a potential target that can hinder the *P. falciparum* growth, as membrane fusion and formation of protein complexes are essential for the parasite’s survival. PF14_0027 codes for a single copy of ubiquitin fused to the ribosomal proteins S27a [14]. Green et al. (2020) found that ubiquitin activation plays an important part in schizont maturation of *P. falciparum* blood-stage regulated by ubiquitin activating enzyme (UBA1) [13]. The study reported the inhibition activity of UBA1 by MLN7243, and can be helpful for the final differentiation of schizonts to merozoites. PF3D7_1434300 is known to be a stress-induced-phosphoprotein 1 (STI1)-like protein that facilitates the assembly of Hsp70/Hsp90 [12]. It has been proposed that these heat-shock proteins with an additional interaction with putative Hop homologue (PfHop) in *P. falciparum* is crucial for the folding and function of proteins involved in signal transduction; thus the inhibition of these protein interactions can induce detrimental effects on the parasite [20]. PF3D7_1116800 heat shock protein 101 (HSP101) plays a crucial role in protein export across the parasitophorous vacuole membrane (PVM), and the blockage of HSP101 leads to the disruption of protein export, causing impaired parasite growth and development [5]. With the wide range of proteins interacting with PF3D7_1438600, if positive correlation is assessed between the interactions, it suggests that the inhibition of PF3D7_1438600 can also hinder the functions of these proteins to induce a more effective and desirable therapeutic response.

**Figure 2.**
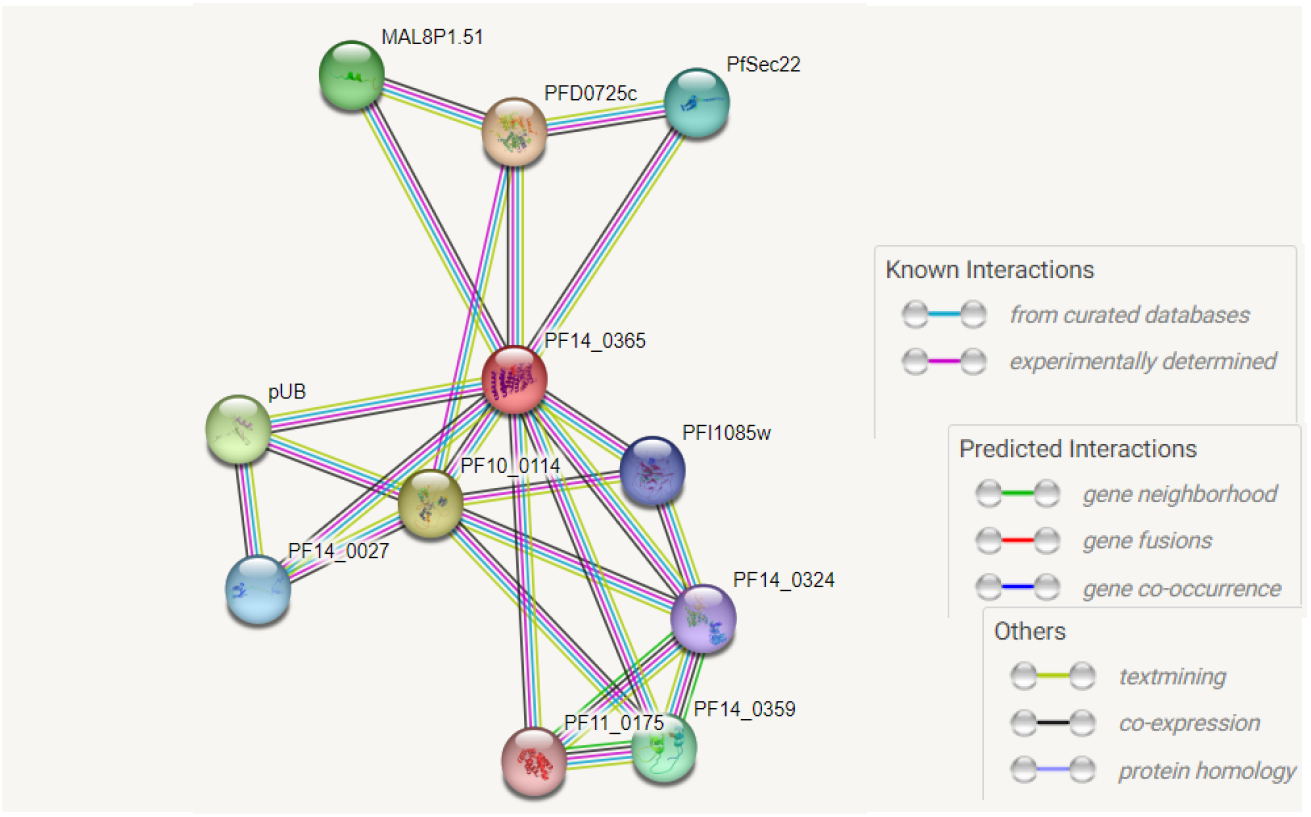
Protein-protein interaction of PF3D7_1438600.

## Conclusions

Novel and suitable drug targets against P. falciparum were identified by utilizing recently published novel proteins and employing proteomics analyses along with in silico druggability approaches. As a result, 7 proteins were identified to have the capability of becoming a potential drug target according to their druggability score. The proteins with the best drug probability score are tied between PF3D7_1455400 and PF3D7_0418400 with both having a score of 1, while the others were still good enough with the lowest score of 0.84 (PF3D7_1204500). Even so, when the protein-protein interaction was taken into account, it was found that only PF3D7_1438600 exhibited strong connections with other *P. falciparum* proteins. Therefore, it can be concluded that PF3D7_1438600 is the most favorable drug target, as it demonstrated a druggability score of 0.91 and the potential to induce disruption when targeted. The identification of such drug targets offers promising opportunities for developing novel strategies to address the challenge of *P. falciparum* drug resistance.

